# High-Throughput Translational Profiling with riboPLATE-seq

**DOI:** 10.1101/819094

**Authors:** Jordan B. Metz, Nicholas J. Hornstein, Sohani Das Sharma, Jeremy Worley, Christian Gonzalez, Peter A. Sims

## Abstract

Protein synthesis is dysregulated in many diseases, but we lack a systems-level picture of how signaling molecules and RNA binding proteins interact with the translational machinery, largely due to technological limitations. Here we present riboPLATE-seq, a scalable method for generating paired libraries of ribosome-associated and total mRNA. As an extension of the PLATE-seq protocol, riboPLATE-seq utilizes barcoded primers for pooled library preparation, but additionally leverages rRNA immunoprecipitation on whole polysomes to measure ribosome association (RA). We compare RA to its analogue in ribosome profiling and RNA sequencing, translation efficiency, and demonstrate both the performance of riboPLATE-seq and its utility in detecting translational alterations induced by specific inhibitors of protein kinases.

## BACKGROUND

The cellular responses to many physiologic stimuli require new programs of protein production. Transcriptional regulation allows direct control of gene expression over a broad dynamic range, but cells can often more rapidly adjust protein levels through translational control. Consequently, alongside transcription factors and their associated regulatory networks, there are mechanisms modulating the translation of specific genes. mTOR is an important example of a translational regulator integrating many potential extracellular signals to regulate cellular metabolism and protein synthesis. Activated through the PI3K signaling axis, mTORC1 phosphorylates eIF4E inhibitors (4E-binding proteins, or 4E-BPs), which releases eIF4E and promotes formation of the eIF4F complex in the initial steps of translational initiation^1^. These effects are mediated by 5’ terminal oligopyrimidine (5’TOP) motifs, which are C/T-rich sequences in the 5’ UTRs of mTORC1 target transcripts^2^. The mTOR protein, the 4E-BP/eIF4E axis, and the 5’TOP motif-containing genes (TOP genes) constitute a basic translational regulatory network.

Despite the attention garnered by profiling transcriptional regulatory networks, less progress has been made in understanding systems-level translational control, in part due to a lack of scalable methods for coupling measurements of protein synthesis with a large number of perturbations. Early studies of translational regulation combined polysome profiling and microarrays to quantify ribosome association genome-wide^3^. The combination of nuclease footprinting of ribosomes^4^ and deep sequencing led to the development of ribosome profiling, which refines translational profiling by resolving the positions of bound ribosomes throughout the transcriptome^5^.

Though amenable to detailed mechanistic analyses in a small number of samples, these approaches are prohibitively expensive and labor-intensive to scale for concurrent analyses across a large sample set. The cost in time and resources for these processes and intervening purifications renders ribosome profiling a suboptimal readout for a broad screen of translational regulation due to lack of scalability. For ideal systems-level analysis of protein synthesis, genome-wide perturbations coupled to a genome-wide readout of translation would allow direct observation of changes in response to a systematic screen of potential perturbations across a large number of conditions and replicates.

Here we present riboPLATE-seq, a scalable method for generating paired libraries of ribosome-associated and total RNA, based on our Pooled Library Amplification for Transcriptome Expression (PLATE-seq) technology^6^. PLATE-seq utilizes sample-specific barcodes, added during reverse transcription, to pool cDNA from multiple individual samples early in library preparation, enabling highly-multiplexed RNA-seq and reducing both reagent and labor costs. Furthermore, as PLATE-seq generates cDNA fragments strictly from the 3’ ends of polyadenylated RNA via oligo(dT)-primed reverse transcription, the resulting libraries are less complex than those with full gene body coverage, and therefore require fewer reads per sample. PLATE-seq consequently has greater throughput than conventional RNA-seq in both library preparation and sequencing.

We utilize riboPLATE-seq for parallel, genome-wide translational profiling in 96-well plates. With PLATE-seq as a genome-wide readout for pan-ribosomal immunoprecipitation, riboPLATE-seq quantifies the ribosome-associated fraction of each polyadenylated transcript relative to total polyadenylated RNA. These paired measurements allow detection of gene-specific changes in ribosome association. While riboPLATE-seq measures the abundance of ribosome-bound mRNA rather than nucleotide-resolution ribosome density as in ribosome profiling, it is highly scalable, inexpensive, and seamlessly compatible with automated liquid handling.

In this study, we use riboPLATE-seq to interrogate translational regulation mediated by the PI3K/mTOR and MAPK/ERK signaling pathways in TS543 glioma neurospheres, which harbor PDGFRA amplification and exhibit constitutive activation of several kinases involved in these pathways^7^. Both pathways influence the formation of the eIF4F complex, composed of the cap-binding eIF4E, the helicase eIF4A, and the scaffold eIF4G. This complex is required in cap-dependent translation initiation, widely regarded as the rate-limiting step in protein synthesis^5,8,9^. In a kinase cascade, PI3K phosphorylates AKT, which phosphorylates mTOR, which then phosphorylates 4E-BPs and releases eIF4E, facilitating eIF4F formation.^1,2^. Separately, the MAPK signaling cascade activates MNK1 which phosphorylates eIF4E directly, increasing its affinity for the 5’ m7G cap and stabilizing eIF4F^10^. The interaction between MNK1 and mTOR is of particular interest in deciphering translational regulation due to their convergence. By disallowing eIF4E release from 4E-BPs, rapamycin indirectly prevents its phosphorylation by MNK1, but multiple human cancer cell lines demonstrate a paradoxical increase in MNK-dependent eIF4E phosphorylation under mTOR inhibition^11,12^. Furthermore, while rapamycin rapidly reduces phospho-eIF4E, MNK1-dependent eIF4E phosphorylation demonstrably occurs under prolonged treatment^13^. MNK1 blockade has additionally been demonstrated to sensitize rapalog-resistant glioma to inhibition of mTORC1^14,15^.

Many previous studies of the effect of MNK1/2 on translation utilize CGP57380^11,14,16,17^, an inhibitor with notable off-target effects^18–22^. CGP57380 has been found to impact initiation and polysome assembly outside the known effects of MNK1 on eIF4F formation, sharing targets with both the known multi-kinase inhibitor imatinib and the specific RSK inhibitor BI-D1870, including inhibition of S6K and 4E-BP1 phosphorylation independent of MNK signaling^19^. CGP57380 was also noted to inhibit several kinases with similar potency to MNK1, including MKK1, BRSK2, and CK1^21^. Remarkably, increased eIF4E:4EBP1 binding, presumed to be a result of MNK1 inhibition and subsequent loss of eIF4E phosphorylation, was demonstrated at CGP57380 concentrations below those inhibiting MNK1^23^. Therefore, the specific function of MNK1 in regulating protein synthesis remains unclear. MNK-i1, recently identified as a highly specific and potent MNK inhibitor with IC_50_ of 0.023 μM and 0.016 μM for MNK1 and MNK2 respectively (compared with 0.87 and 1.6uM for CGP57380), blocks eIF4E phosphorylation without impacting other pathways converging on eIF4E^23^. Thus, we sought to clarify the effect of MNK1 with this novel inhibitor.

We demonstrate the riboPLATE-seq technique in a screen of translational regulators downstream of amplified PDGFRA signaling in TS543 cells, including kinases mTOR, PI3K, and MNK1, and additionally generate signatures of mTOR inhibition with ribosome profiling and RNA sequencing for comparison to established methods of interrogating translation. Importantly, ribosome profiling provides a measure of ribosome-mRNA association that is quantitatively distinct from riboPLATE-seq. The ratio of normalized aligned reads in ribosome profiling over RNA-seq libraries obtained from the same biological sample, referred to as translation efficiency (TE), corresponds to the average number of ribosomes bound per transcript. In contrast, the ratio of normalized aligned reads in riboPLATE-seq over normal PLATE-seq relates to the fraction of the available pool of transcripts bound by ribosomes, which we term ribosome association (RA). Despite their differences, we expect significant overlap in the effects revealed by these measurements, especially in response to inhibition of kinases involved in translational regulation.

## RESULTS

### Overview of riboPLATE-seq Technology

riboPLATE-seq enables transcriptome-wide measurements of ribosome association in a multi-well plate by combining pan-ribosomal immunoprecipitation (IP) with a low-cost technique for RNA sequencing, PLATE-seq (Figure 1a). In PLATE-seq, we isolate polyadenylated transcripts with an oligo-(dT) capture plate, followed by incorporation of well-specific barcodes in poly(dT)- primed reverse transcription. After pooling barcoded cDNA libraries across each plate, we conduct all subsequent library preparation steps on the pool and sequence the resulting libraries to a modest depth (few million reads per well). Previous studies isolate ribosome-bound mRNA from specific cell types *in vivo* with the translating ribosome affinity purification (TRAP)^24^ and RiboTag^25^ systems, relying on transgenic or recombination-driven epitope labeling of ribosomal proteins for immunoprecipitation. We instead use a native epitope in 5.8S rRNA for pan-ribosomal IP. By comparing transcript abundance measured by PLATE-seq with and without ribosomal IP for all genes, we measure gene-specific ribosome association for all genes detected, extending the scalability of PLATE-seq from transcriptional to translational profiling.

**Figure 1:**
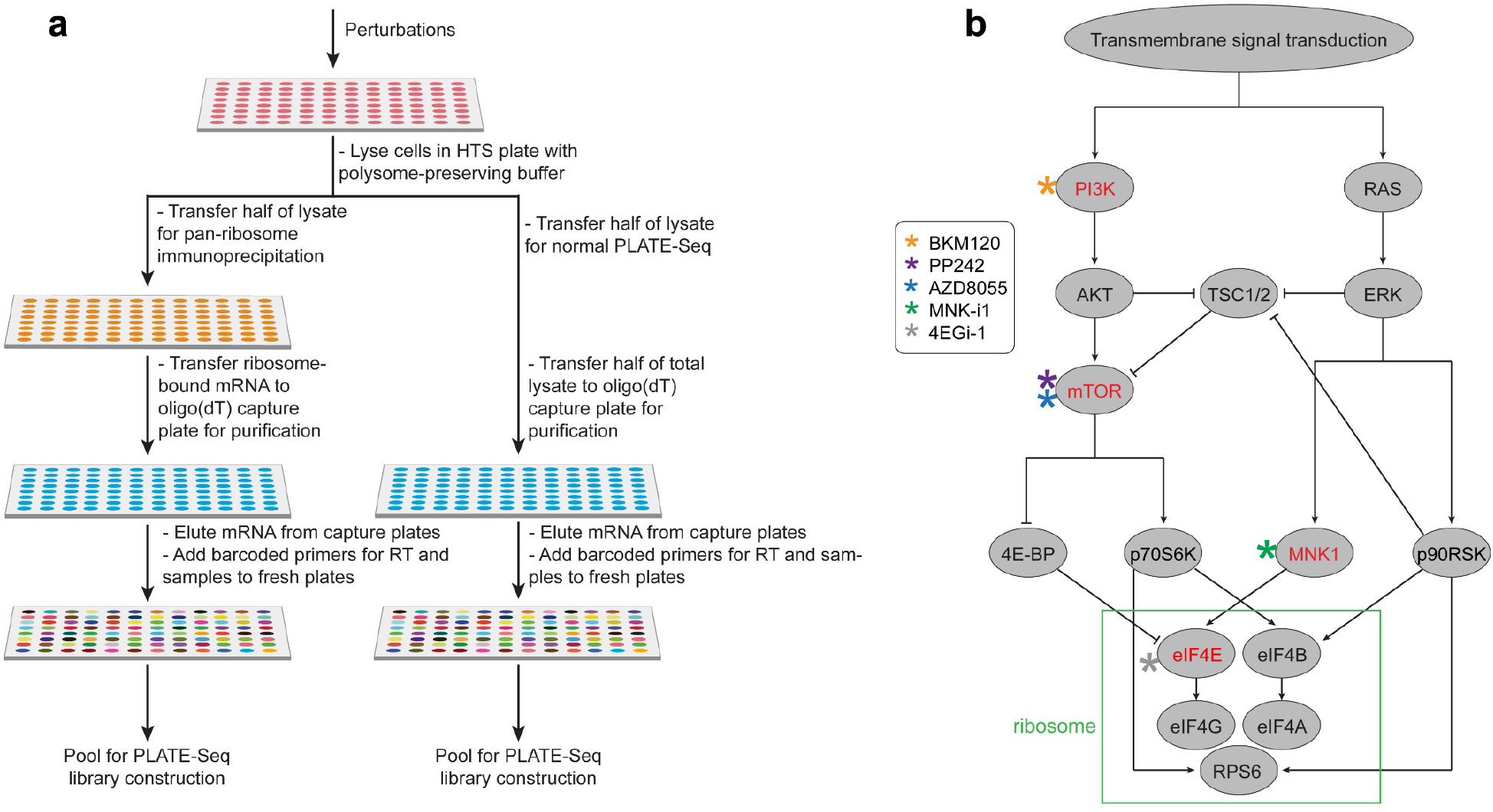
Overviews of the protocol and experimental design of the study performed. **a)** Schematic diagram of the riboPLATE-seq protocol, from lysis in a multi-well plate to pooled library preparation. The right-hand side mirrors the original PLATE-seq protocol. In this workflow, an oligo(dT)-grafted plate captures polyadenylated RNA that can be reverse-transcribed with barcoded adapters, generating a plate of cDNA that may be pooled for library construction. The left side incorporates a pan-ribosome IP before PLATE-seq pooling and library preparation, generating instead a pooled library of ribosome-associated RNA. **b)** Simplified structure of the signaling pathways under study and the specific protein targets considered. The PI3K/AKT/mTOR signaling axis at left converges with the MAPK/ERK pathway at right on eIF4E, early in the process of ribosome assembly (green box). The figure also outlines the inhibitors used in this study and their specific targets within these pathways. NVP-BKM120 is a PI3K inhibitor (orange), both AZD-8055 and PP242 are mTOR inhibitors (blue and purple, respectively), MNK-i1 is a MNK1 inhibitor (green), and 4EGi-1 is a direct eIF4E inhibitor (black).

To implement riboPLATE-seq, we divide polysome lysates from a multi-well plate into two plates. We subject one plate to indirect, pan-ribosomal IP with robotic liquid handling, using biotinylated anti-rRNA antibody y10b and streptavidin-coated magnetic beads. Finally, we generate PLATE-seq libraries from the immunoprecipitated polysomes in one plate and total lysate from the other, as described previously. This design minimizes sample-to-sample noise by processing up to 96 samples in one batch with automated liquid handling, which further reduces the time and effort required to process large numbers of samples. Per-sample reagent costs are also substantially lower, as PLATE-seq generates 3’-end RNA libraries approximately $12 per sample, while riboPLATE-seq requires additional expenses for automated ribosome IP totaling ~$13 per sample for reagents and disposables, all in 96-well plates. A full riboPLATE-seq study performed on plate costs ~$25 in materials per sample to generate paired libraries of ribosome-associated and total RNA from cellular lysates, compared with $122 for ligation-free ribosome profiling and RNA sequencing.

### riboPLATE-seq Translational Profiling Experimental Overview

To demonstrate the utility of riboPLATE-seq, we measured the translational impact of inhibiting different components of signaling pathways in TS543 that converge on the ribosome (Figure 1b). We sought to identify targets specific to each regulator and compare translational signatures by screening several regulators in these pathways. The drug treatments consisted of two competitive mTOR inhibitors, PP242 and AZD-8055; an inhibitor of PI3K, BKM120; a specific inhibitor of MNK1/2 activity, MNK-i1; and 4EGi-1, a 4E-BP mimic that inhibits the association of eIF4E and eIF4G. We determined concentrations of these drugs from an examination of the literature, ensuring values near the half-maximum inhibitory concentrations (IC_50_) for the main substrates of the drugs in question: 625nM PP242^26^, 50nM AZD-8055^27^, 1μM BKM120^28^, 100nM MNK-i1^29^, and 50μM 4EGi-1^30^. In order to analyze interactions between kinases, we also treated samples with pairwise combinations of PP242, BKM120, and MNK-i1 at the same concentrations. For comparison, we treated TS-543 neurospheres with PP242 at the same concentration as in the riboPLATE-seq experiment at multiple time points to assess the similarity of translational perturbations detected across experiment types. We expected these two methods to give quantitatively distinct results while identifying similar target sets. An overview of all sequencing data generated for this study is provided in Supplementary Table S1.

### Immunoprecipitation of Ribosome-bound mRNA

To assess the depletion of free RNA in the riboPLATE sequencing libraries following pan-ribosome immunoprecipitation, we added ERCC spike-in to half of the wells in a riboPLATE-seq experiment without automated liquid handling, after lysis but prior to ribosome IP. Figure 2a shows the distributions of total spike-in RA and total transcript RA across wells. Spike-ins exhibit almost uniformly lower RA than transcripts, with total RA across wells of 0.25 and 1.04, respectively (one-tailed paired-sample Wilcoxon signed-rank p=1×10^−9^). The relative depletion of spike-ins to genes for each well shows depletion across the set spike-ins relative to the transcriptome in all but two wells (2/48), which demonstrated modest (<0.5 log_2_-fold) enrichment (Figure 2b).

**Figure 2:**
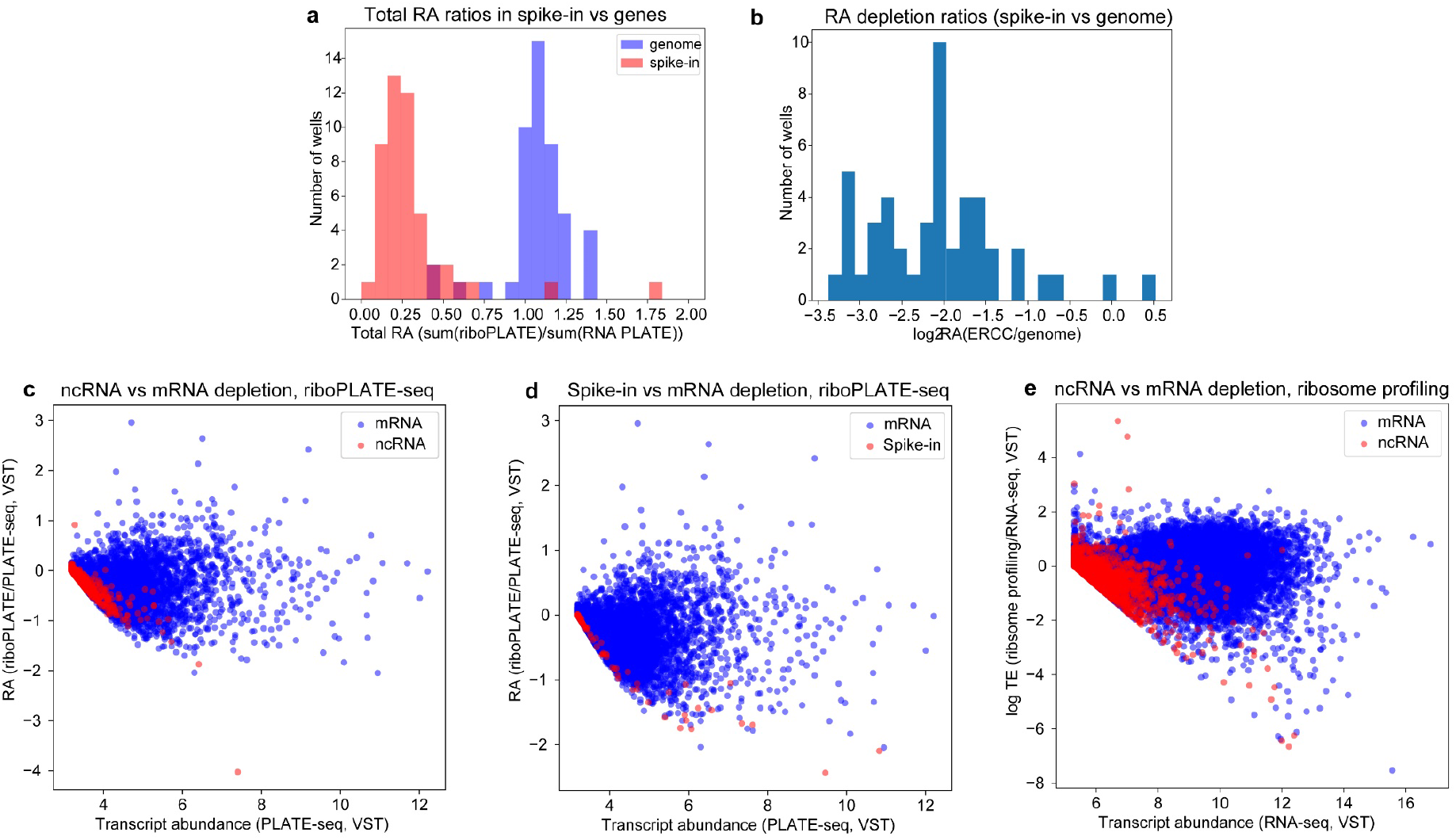
Assessment of riboPLATE-seq IP fidelity. **a)** Depletion calculated per-sample as the log2-ratio of the sum of all spike-in or gene-aligned counts in the riboPLATE-seq library over the same sum in the sample’s paired PLATE-seq library, using DESeq2-normalized counts (median of ratios normalization). Spike-ins show more significant depletion than genes in almost all wells (mean spike-in RA 0.25; mean genomic RA 1.04; Wilcoxon signed-rank test p=1×10^−9^) **b)** The same information in **a**, presented as the per-well difference in depletion ratios for ERCC and the background genome, demonstrating significant depletion of spike-in RNA with IP in most libraries (mean log2 depletion ratio −2.1). **c)** Relationship between transcript abundance (in PLATE-seq) and RA (riboPLATE/PLATE-seq) for coding genes and non-coding RNA (ncRNA). ncRNA are heavily depleted (RA<0) at all expression levels, to a greater extent than almost all genes. **d)** Relationship between transcript abundance and RA for ERCC spike-ins vs coding genes, demonstrating a pronounced pattern of depletion in RA for spike-ins relative to mRNA across all expression levels. **e)** The corresponding plot to **c** derived from ribosome profiling and RNA sequencing data of TE and RNA-seq transcript abundance. Though the shape of the distribution is different, ncRNA still demonstrate lower TE than mRNA at higher expression levels.

As ribosome profiling has revealed low but significant levels of ribosome occupancy among ncRNA^31^, we sought to contrast ribosome association between ncRNA and mRNA with riboPLATE-seq, expecting noncoding transcripts to be depleted from riboPLATE-seq relative to PLATE-seq. Examination of the relationship between RA and underlying transcript abundance reveals lower RA for ncRNA than mRNA at all expression levels (Figure 2c), with similar distribution for ERCC spike-ins in the riboPLATE-seq samples containing them (Figure 2d). Depletion is also observed for ncRNA in ribosome profiling and RNA sequencing, with lower TE than mRNA on average over all RNA-seq expression levels (Fig. 2e). Combined with the observed spike-in depletion, our results are consistent with depletion of free RNA by the pan-ribosomal immunoprecipitation implemented in riboPLATE-seq.

### Assessment of riboPLATE-seq Library Quality

We sought to establish the quality of the pooled ribosome-associated and normal PLATE-seq libraries with regard to read depth, library complexity, and saturation. Figures 3a-b show gene detection saturation curves for riboPLATE-seq and PLATE-seq, demonstrating the dependence of these libraries’ sensitivities on read depth, while Figure 3c demonstrates their range in complexity at full depth. Read depth and library complexity were similarly distributed in riboPLATE- and PLATE-seq libraries in our kinase screen with automated liquid handling. We detected ~9-11K unique genes at full depth in either library type, with riboPLATE-seq detecting slightly fewer genes in about twice as many reads at saturating sequencing depth.

**Figure 3:**
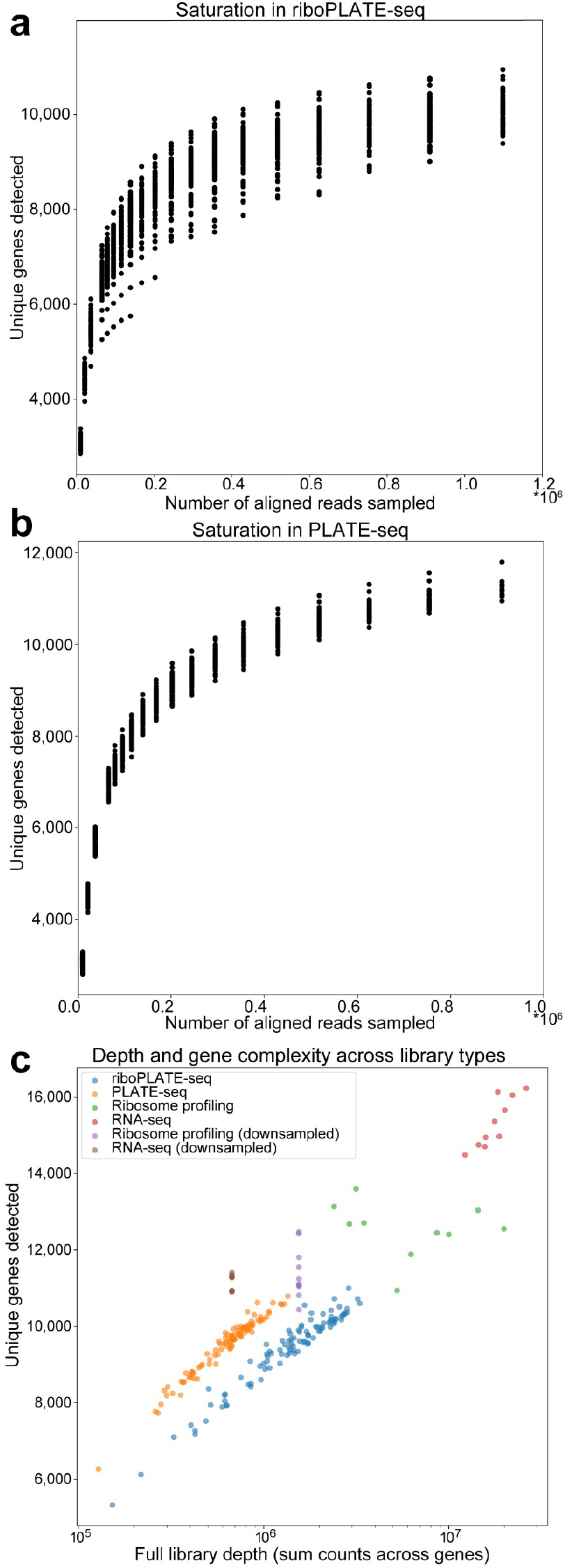
Analysis of riboPLATE-seq library saturation, size, and complexity. **a,b)** Library saturation strip plots for ribosome-associated (riboPLATE-seq) and total RNA (PLATE-seq) libraries in this study. In each, the Y axis shows the number of unique genes detected in each sample at each subsampled read depth on the X axis, excluding libraries smaller than the subsampling depth. With ~10-11,000 unique genes detected, riboPLATE-seq and PLATE-seq are comparably saturated. **c)** Scatter plots emphasizing the relationship between library size and complexity across library types. The Y axis represents the number of unique genes detected within a library; the X axis represents its size in summed gene counts. PLATE-seq and riboPLATE-seq are very similarly distributed, with PLATE-seq generating slightly more complex, smaller libraries than riboPLATE-seq. Ribosome profiling and RNA-seq generate larger, more complex libraries than either riboPLATE- or PLATE-seq, which retain their complexity with ~11,000 genes detected when downsampled to the average read depths of riboPLATE- and PLATE-seq, respectively.

Figure 3c highlights the differences between riboPLATE-seq and PLATE-seq in terms of library complexity and sequencing depth, additionally contrasting with full-length ribosome profiling and RNA-seq. On average, automated riboPLATE-seq detects approximately 9,940 unique genes in 1.56 million uniquely mapped reads per sample, while PLATE-seq detects an average of 10,710 genes in 0.67 million (677K) reads per sample in this study. These measurements are comparable to the initial report characterizing PLATE-seq, which detected an average of approximately 10,200 genes from 0.67 million uniquely mapped reads^6^. In contrast, ribosome profiling and total RNA sequencing libraries detect an average of 15,000 and 15,900 genes respectively at full sequencing depth, and 12,900/12,300 genes each when downsampled to the respective median read depths of riboPLATE-seq and PLATE-seq libraries, reflecting their inherent complexity even at lower depth. In summary, riboPLATE-seq preparation following pan-ribosome IP generates libraries that are comparable to their unmodified PLATE-seq counterparts in terms of saturation and complexity.

### Pharmacological Kinase Screen with riboPLATE-seq

After establishing the performance of riboPLATE-seq, we sought to characterize its ability to detect changes in RA in a screen of signaling pathways converging on the ribosome. We first calculated RA for each gene in each sample and the change in RA for each drug-treated sample relative to the mean across vehicle-treated controls, generating individual signatures of absolute RA and log-fold change in RA (lfcRA) for every sample relative to control. Principal component analyses (PCA) of RA and lfcRA both reveal separation of samples according to their RA change relative to control in the first component (PC1), clustering samples into condition-specific groups in Figures 4a-b. In Figure 4b, the mean PC1 value across samples in a condition is inversely correlated with the number of genes exhibiting significant differences in RA for that condition relative to vehicle-treated controls (Figure 4c, Spearman r= −0.90, p=0.02). PC1 values for each individual sample are also inversely correlated with mean effect size, calculated as the average of absolute log-fold changes in RA across genes determined significant in any condition by DESeq2 (Spearman r=−0.74, p=6.9×10^−8^). PC2 further separates most samples treated with drug combinations or 4EGi-1 from those treated with individual kinase inhibitors, such that combination-treated clusters aggregate more closely with each other than with any of their individually-treated counterparts.

**Figure 4:**
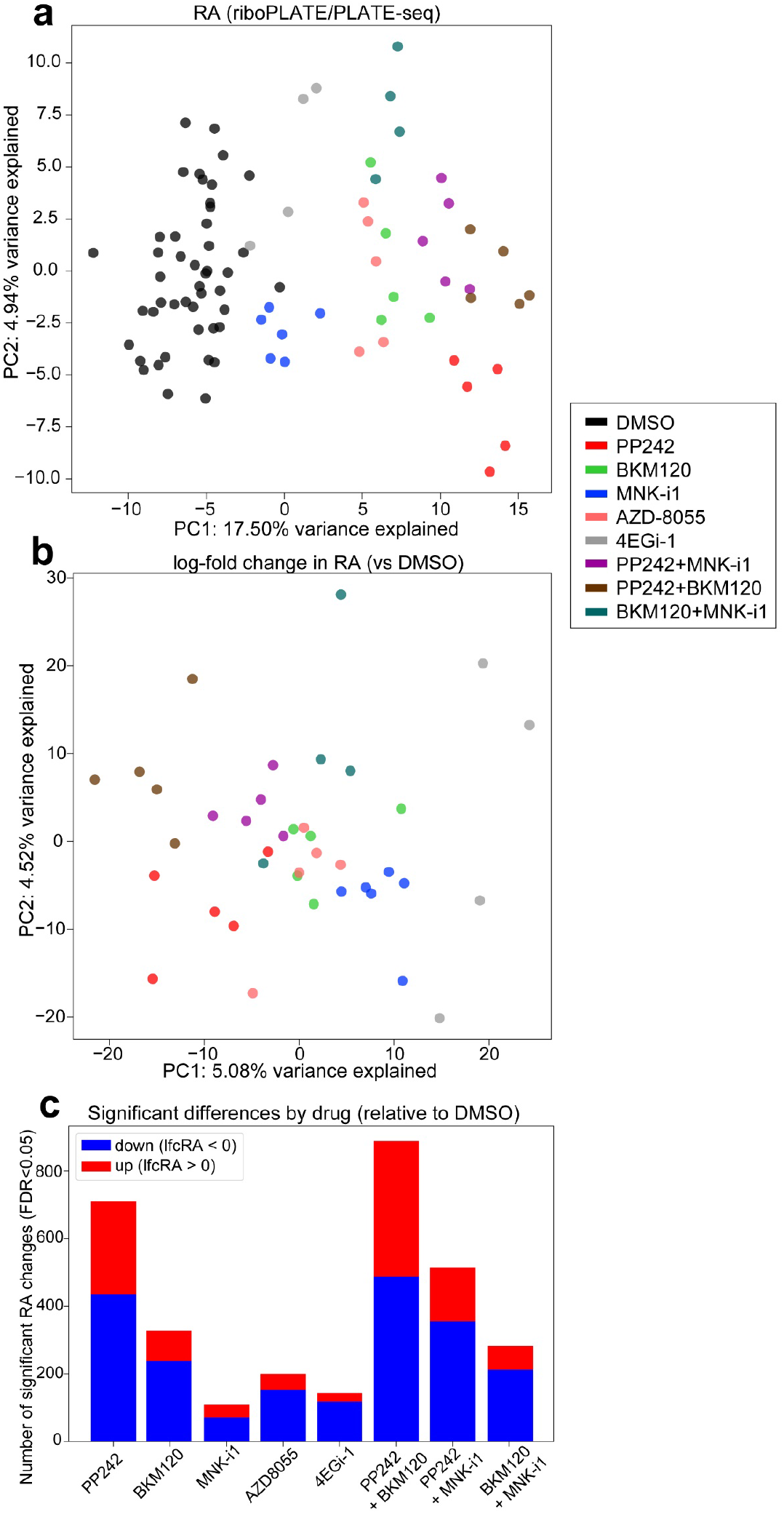
Principal Component Analyses (PCA) of riboPLATE-seq data. **a)** riboPLATE-seq ribosome association (RA, riboPLATE-seq-PLATE-seq), **b)** difference in RA between each sample and the average across DMSO-treated samples (lfcRA), normalized by variance-stabilizing transform (VST) in DESeq2. For both plots, the domain of the PCA was restricted to genes with significant changes in RA reported by DESeq2 for any drug treatment relative to DMSO, (FDR<0.05, 1,813 genes total; detailed in Fig. 4c). Drug treatments elicit changes consistent enough to yield clustering behavior among samples treated with the same drug in both analyses, as well as co-clustering of related drug treatments (e.g. BKM120, PP242, and AZD8055). Separation is also apparent between combination treatments and their constituent, individual drugs in each plot. **c)** Significant effects of each drug determined in riboPLATE-seq by DESeq2 (Benjamini-Hochberg adjusted false discovery rate (FDR) < 0.05). Genes determined significantly up- or downregulated in ribosome association (RA) are tallied for each drug and combination treatment.

In both plots, clusters of samples treated with singular kinase inhibitors are organized consistently with their expected effects. Samples corresponding to mTOR axis inhibitors (PP242, BKM120, and AZD8055) cluster near each other, while MNK-i1-treated samples cluster more closely with DMSO-treated controls in Fig. 4a and 4EGi-1-treated samples in Fig. 4b. Together with the relatively few genes exhibiting significantly changed RA due to MNK-i1 treatment in Fig. 4c, this suggests that MNK-i1 has a weaker impact on ribosome association than any drug or combination tested.

### Differential Ribosome Association and Translation Efficiency: Comparison of riboPLATE-seq and Ribosome Profiling

In order to more rigorously analyze differential RA as a function of drug treatment, we used DESeq2 to compare each drug to control with riboPLATE-seq. We separately calculated differential translation efficiency from ribosome profiling and RNA sequencing data from PP242-treated and control samples, for comparison with established translational measurements. Figures 5a-e display the volcano plots of log-fold change in RA (lfcRA) vs statistical significance (−log_10_(FDR), which is inversely proportional to Benjamini-Hochberg adjusted false-discovery rate) for all genes in each individual drug treatment tested. Figures 5f,g display the analogous plots for differential TE from ribosome profiling under 30-minute and 6-hour PP242 treatment, overlaid with the genes significantly changed under PP242 treatment in riboPLATE-seq (FDR<0.05, red/blue circles)emphasizing the correlation of effects across methods. PP242 targets determined by significant lfcRA exhibit similar changes in TE, with upregulated targets demonstrating significantly higher TE than downregulated targets at both 30 minutes and 6 hours of PP242 treatment (one-tailed, two-sample Mann Whitney U test p=3.0×10^−64^ and p=1.7×10^−92^, respectively). In particular, we observe significant downregulation of TOP motif-containing genes in both riboPLATE-seq and ribosome profiling, marked green in all plots. This set of canonical mTORC1 targets exhibits significantly reduced TE after both 30 minutes and 6 hours of treatment with PP242 (one-tailed, two-sample Mann-Whitney U test p=1.2×10^−38^ and p=2.9×10^−46^, respectively), with likewise significant reductions in RA associated with mTOR axis inhibitors (PP242, AZD8055, and BKM120, p= 4.0×10^−26^, 8.9×10^−19^, and 1.6×10^−14^, respectively). Though similarly reduced in MNK-i1 and 4EGi-1 treatment, these reductions were smaller and less significant (p=0.00002 and p=0.86, respectively). Both drugs also induced fewer statistically significant changes in RA than the other drugs tested (Figure 4c). While for MNK-i1 this might be attributed to a weaker overall effect for the drug, 4EGi-1 elicits many high-magnitude changes in RA that do not achieve statistical significance.

**Figure 5:**
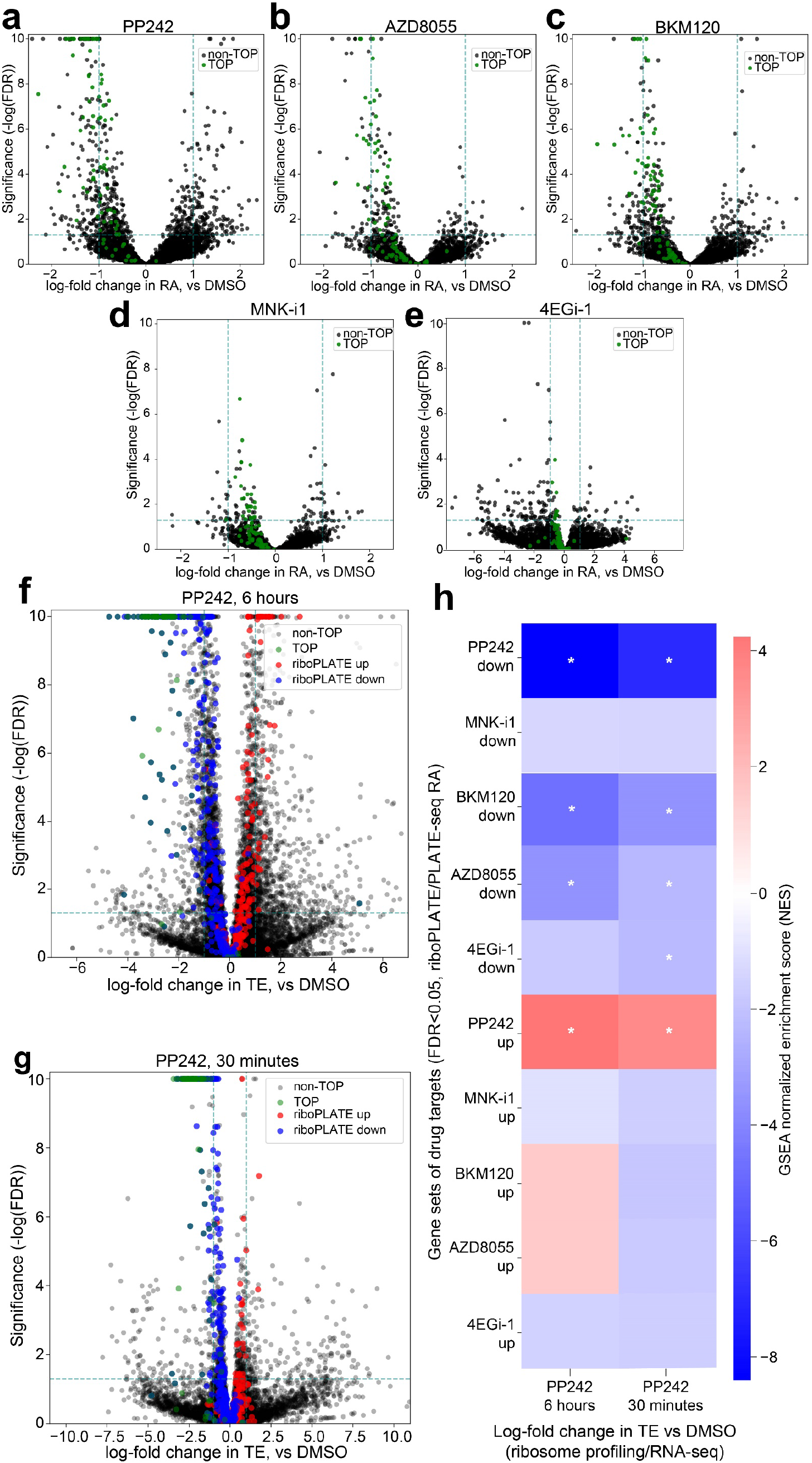
Characterization of Alterations in Ribosome Association (RA) and Comparison with Translation Efficiency (TE) from Ribosome Profiling and RNA Sequencing. **a-e)** Volcano plots of log-fold change in RA per-gene associated with each drug treatment. In each plot, the X axis marks the size and direction of the observed change for each gene, while the Y axis scales inversely with p-value (−log_10_(p)) to provide a positive metric of significance; TOP motif-containing genes, the canonical targets of mTOR signaling, are colored red. Additionally, dashed lines at y=−log_10_(0.05) and x = +/− 1.0 provide an estimate of the number and magnitude of significantly up- and down-regulated genes for each condition. Significant TOP gene inhibition is seen in treatment with PP242, AZD8055, and BKM120, but less so with MNK-i1 and not at all with 4EGi-1. **f,g)** Volcano plots of log-fold change in TE per-gene associated with PP242 treatment for 30 minutes or 6 hours, generated from ribosome profiling and RNA sequencing data. Guide lines at y=−log_10_(0.05) and x=+/−1 provide gauges of the significance and magnitude of the effects on gene-wise TE at either time point. TOP motif-containing genes, in red, are highly and significantly downregulated in both (Mann-Whitney U test p=2.9×10^−46^ and p=1.2×10^−38^ for 6-hour and 30-minute treatment, respectively). In all volcano plots, genes with p-values less than p=1×10^−10^ (y=10) are given a maximum y-value of 10 to prevent skewing of the axes. **h)** Enrichment heatmap using GSEA to compare signatures of differential TE (X axis) with sets of genes significantly up- or downregulated (DESeq2 FDR<0.05) at the level of RA by different 6-hour drug treatments determined with riboPLATE-seq. Genes affected by PP242, BKM120, and AZD8055 are concordantly altered in TE after 6 hours of PP242 treatment, while only the genes impacted by PP242 at the level of RA are similarly affected in TE after 30 minutes of PP242 treatment, with all other gene sets demonstrating varying degrees of downregulation (GSEA Normalized Enrichment Score < 0).

We further compared riboPLATE-seq and ribosome profiling using Gene Set Enrichment Analysis (GSEA)^32^, focused on the sets of genes determined significantly upregulated or downregulated in RA by DESeq2 for each condition (FDR<0.05). We first made these sets of genes exclusive to each other, such that no two downregulated nor upregulated sets shared a common gene, then observed their distribution in log-fold change in TE (lfcTE) following PP242 treatment with the pre-ranked function in GSEA. Figure 5h describes the results of this analysis as a heatmap, overlaid with asterisks (*) for gene sets determined significantly up-or downregulated in lfcTE (Bonferroni-adjusted FWER<0.05). Genes corresponding to riboPLATE-seq PP242 targets demonstrate expected, concordant changes in lfcTE, with upregulated targets enriched for increased TE and downregulated targets depleted. The targets for BKM120 and AZD8055 are similarly distributed at 6 hours of PP242 treatment, to lesser degrees of significance, while they are uniformly weakly downregulated at 30 minutes. The targets of 4EGi-1 and MNK-i1 are also downregulated in TE at both time points, regardless of their direction of RA change in riboPLATE-seq. Our results demonstrate the capability for riboPLATE-seq to detect specific translational changes that are reflected in ribosome profiling, despite the quantitative differences in the measurements of RA and TE.

### Attenuation of Perturbations to Ribosome Association in Drug Combinations

While we expected some degree of synergy in the effects of individual drugs in combination treatments, or at least additivity of effects from simultaneous inhibition of both the PI3K/mTOR and MAPK/ERK pathways, we instead found that the strongest effects of individual drugs were attenuated when combined. Fig. 6a-c are volcano plots of differential RA for each drug combination, color-coded to indicate up- and downregulated targets of the combination’s respective constituent drugs determined via DESeq2 (FDR <0.05). In all plots, individual targets demonstrate reduced significance under combinations despite the existence of many significant RA changes. We directly compare the magnitude of effect sizes observed for significant effects of the individual drugs in individual and combination drug treatments in Figures 6d-f, finding targets of either individual drug to be largely attenuated in their effects in combination treatments (below the dashed y=x line indicating equality of effect in both conditions). We find 78-89% and 86-95% of significant downregulated and upregulated targets, respectively, for both individual drugs to be attenuated in their effect in combination treatments, with the combination PP242+BKM120 diminishing the fewest of its individual drugs’ effects. Uniformly across combinations, we found more attenuation in upregulated than downregulated targets of individual drugs, with greater variation in the range of their observed effects in combination treatments. This suggests poor conservation of upregulatory effects in combination treatments relative to their individually-downregulated counterparts. The surprising interactions between drugs in combination treatments raise the possibility of accelerated compensatory responses to combinations of individual inhibitors, with attenuation of individual drugs’ effects corresponding to pronounced compensation in the context of greater second-order, indirect effects. These findings underscore the value of riboPLATE-seq in screening a broad set of potential translational perturbations, allowing direct comparison of individual and combination effects with simultaneously-generated data.

**Figure 6:**
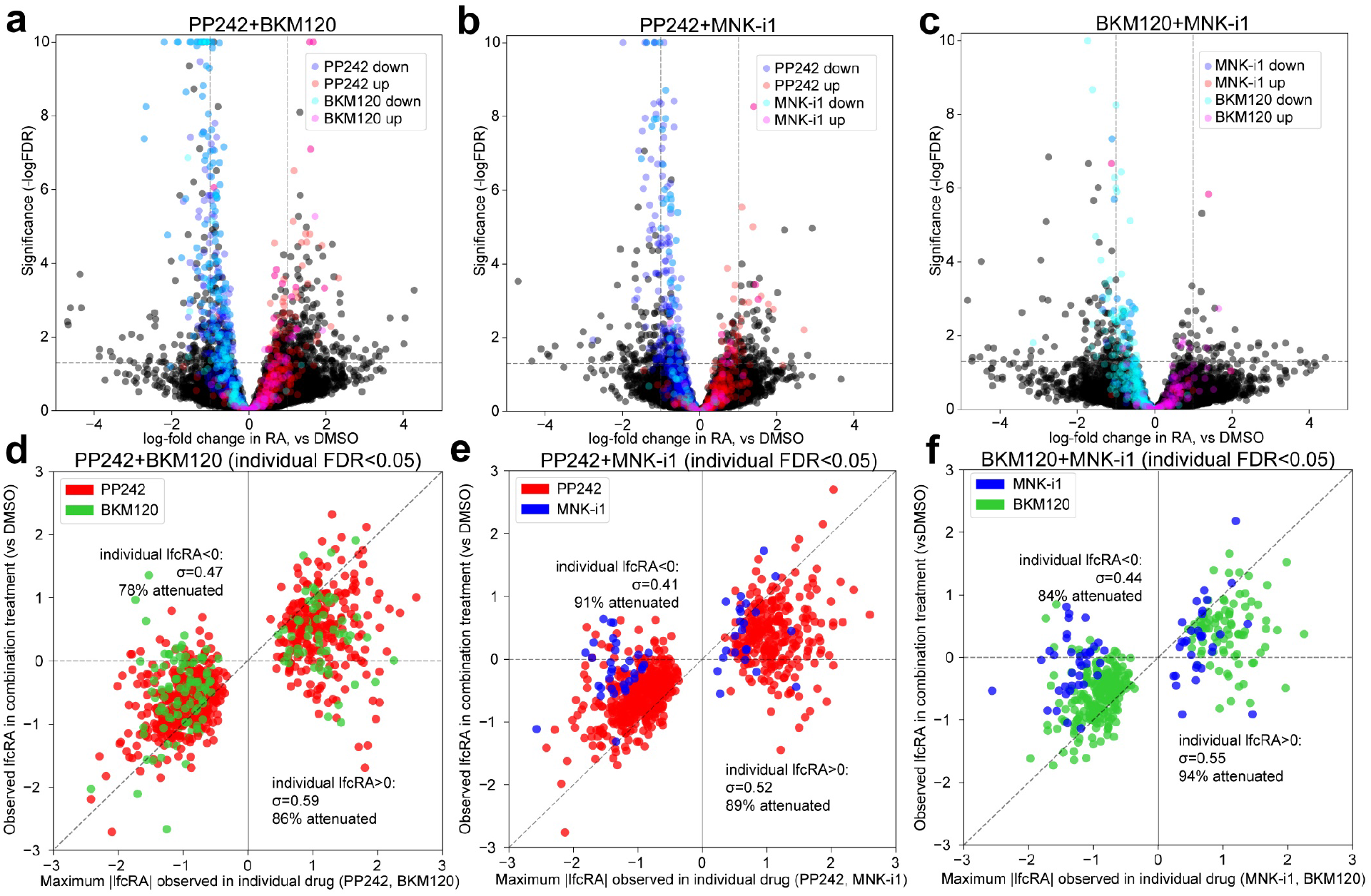
Attenuation of Single-Drug Effects Under Combination Treatments. **a-c)** Volcano plots of changes in RA observed under drug combinations, color-coded with the upregulated (red, magenta) and downregulated (blue, cyan) significant targets (DESeq2 FDR<0.05) of each combination’s constituent drugs. In each plot, the X axis marks each gene’s observed difference in RA under drug combination treatment relative to DMSO controls, and the Y axis scales inversely with p value (−log_10_(FDR)) to correlate positively with the effect’s significance. Significant targets of both constituent drugs are attenuated in their effects in all combinations, indicated by significance values below the guideline y=−log_10_(0.05) and effect sizes reduced towards x=0, relative to their distributions in the volcano plots in Fig. 5 (by definition y>−log_10_ (0.05)). **d-f)** Scatterplots comparing the maximum observed effect in either individual drug on the X axis with its observed effect under combination treatment on the Y axis, for the set of genes significantly impacted by each drug combination’s two constituent drug treatments. Guidelines at Y=0 and X=Y aid visualization of the attenuation in gene-specific effects of the individual drugs in combination. Most genes fall between Y=0 and X=Y, indicating a lesser change in RA in the same direction under combination treatment compared with its maximum effect observed in singular treatments.

### Mapping the Network Controlling TOP gene Translation

To better visualize the interactions between translation-regulatory kinases, we constructed a translation control network from our results with focus on the TOP motif-containing genes.

Following the observation of TOP gene regulation in several experimental conditions, we sought to compare the structure of our network with the known organization of kinases impacting TOP gene translation. In addition to the set of 97 canonical genes, we obtained a set of 182 candidate TOP motif-containing genes that have homologues in the mouse genome with TOP motifs, identified in a genome-wide screen of transcription start sites^33^. We found these TOP candidates to behave similarly to the canonical set (Fig. 7a). TOP genes and candidates are uniformly downregulated by PP242, BKM120, and AZD8055, though a lesser fraction of candidates exhibit significant effects than canonical targets. No candidates exhibited significant changes in RA under MNK-i1 treatment, while only four were downregulated under 4EGi-1, consistent with their relatively small number of canonical TOP gene targets and significant targets overall.

**Figure 7:**
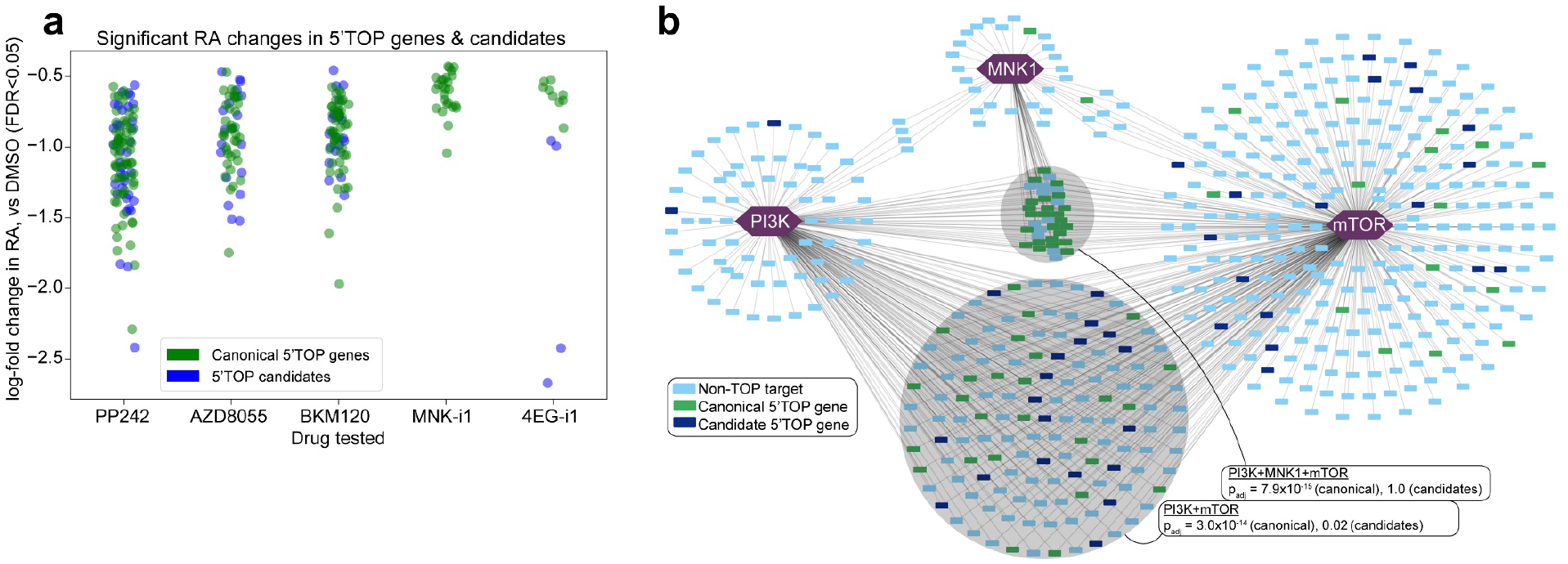
Distribution of TOP Motif-Containing Genes and Candidates in a Modeled Translational Control Network. **a)** TOP genes and candidates significantly perturbed in RA by drug treatments. Strip plots along the X axis, labeled for each drug treatment in our riboPLATE-seq study, contain log-fold changes in RA (Y axis) for the genes exhibiting significant RA perturbations (FDR<0.05) under each treatment relative to DMSO controls, excluding non-TOP-containing genes. TOP candidates behave similarly to canonical TOP genes, exhibiting decreased RA under treatment with mTOR axis inhibitors (PP242, AZD8055, BKM120) while MNK-i1 and 4EGi-1 elicit fewer significant alterations in these sets. **b)** Network representation of targets of mTOR, PI3K, and MNK1, interpreted as the genes exhibiting significant decreases in RA under treatment with PP242, BKM120, and MNK-i1, respectively (FDR<0.05). Targets are color-coded for identification as canonical 5’TOP motif-containing genes (green), TOP gene candidates with known mouse homologues (navy blue) and other genes (light blue). Shaded circles denote subsets of the network with significant enrichment of canonical and/or candidate TOP genes, with the Fisher’s exact test p-value for either set displayed. P-values are corrected for multiple testing over all gene sets and network intersections tested via Benjamini-Hochberg (false discovery rate/FDR) adjustment. The intersection of all three kinases is enriched for canonical TOP genes, but not candidates (Fisher’s exact test padj=7.9×10-15 and padj=1.0, respectively), while the intersection of PI3K and mTOR is significantly enriched for canonical TOP genes and less-significantly enriched for candidates (p_adj_=3.0×10^−14^ and p_adj_=0.02).

A simple translational regulatory network constructed from our riboPLATE-seq data and overlaid with canonical and candidate TOP genes is presented in Figure 7b. We considered the genes demonstrating significant RA reduction under PP242, MNK-i1, and BKM120 as the positive translational targets of mTOR, MNK1, and PI3K, respectively (FDR<0.05). All but one canonical TOP gene and two candidates present in the network are mTOR targets, with significant enrichment of canonical TOP genes and marginally-significant enrichment of candidates in the intersection of targets common to mTOR and PI3K (FDR-adjusted Fisher’s exact test p_adj_=3.0×10^−14^ and p=0.02, respectively), but enrichment only of canonical genes in the intersection of all three kinases (p_adj_=7.9×10^−−15^ and p_adj_=1.0 for canonical and candidate TOP genes, respectively). TOP genes and candidates in the network outside these intersections lie mainly in the exclusive targets of mTOR, though without significant enrichment.

The modeled network is consistent with the known architecture of TOP gene regulation. The strong effects of mTOR and PI3K on TOP gene RA, the relatively minor effect of MNK1, and the observation that the effect of PI3K with respect to these genes is largely a subset of the more direct effect of mTOR can be explained by the known activation of TOP gene translation by mTOR, the activation of mTOR by PI3K, and the minimal demonstrated effect of MNK1 outside this axis^34^ by studies with the more specific MNK-i1 inhibitor^35^.

## DISCUSSION

Ribosome association is frequently used to infer translational activity, whether measured by sucrose gradient fractionation of intact RNA in polysome profiling, or isolation of digested monosomes and their ribosome-protected footprints in ribosome profiling. In polysome profiling, the ratio of transcripts sedimenting in “heavy” vs “light” fractions is similar to RA as defined for riboPLATE-seq. Ribosome profiling refines this measurement with its focus on ribosome footprints, calculating a per-transcript ribosome occupancy with additional information about positions and ribosome arrest^36^. In riboPLATE-seq, we trade this information for increased throughput in library preparation and sequencing to measure ribosome association across the pool of expressed transcripts.

With pooled library construction, greater throughput is possible with riboPLATE-seq than with either ribosome profiling or polysome profiling, though not without limitations. Transcripts with similarity to the y10b epitope may non-specifically bind to the antibody or be adsorbed to the streptavidin-coated magnetic beads, yielding inflated riboPLATE-seq counts, which might be addressed by pre-incubation of antibody-bound beads with exogenous RNA to saturate non-specific binding sites, but care must be taken to avoid ribosomal RNA which could drastically reduce sensitivity. Potential re-initiation in lysate might be minimized with the non-hydrolyzable GTP analogue, GDPNP. As GTP hydrolysis is required in both start site selection and subunit joining steps of 80S initiation complex formation, GDPNP could prevent re-initiation of free ribosomes^37^. Additionally, the method cannot resolve location-specific effects, such as the effect of ribosome association in 5’ leader sequences on translation in downstream coding sequences^38^, and is generally insensitive to translational regulation at the level of elongation as it cannot distinguish active from stalled ribosomes. However, this limitation is common to many measurements of translational activity, including ribosome profiling^39^, and frequently requires more specific analyses of ribosome density^36,40^. In this study, we demonstrate the sensitivity of riboPLATE-seq to perturbations in translation initiation with confirmation of the observed effects in ribosome profiling and RNA sequencing. With a focus on initiation as the rate-limiting step of translation, we expect that the protocol can reveal important regulatory relationships at this level.

We used riboPLATE-seq in this study to interrogate translational regulation in mitogenic signaling pathways in cancer cells, observing the expected effects of mTOR inhibition on translation, including decreased ribosome association in TOP motif-containing genes, and correlated this effect with that seen in differential translation efficiency generated from ribosome profiling and RNA sequencing. We additionally clarified translational targets for PI3K and MNK1, observing that PI3K targets a subset of the TOP genes impacted by mTOR, with no strong independent impact on known TOP genes or candidates. This suggests the effect of PI3K on the TOP genes may be wholly mediated by mTOR, consistent with the known organization of the mTOR signaling axis. In contrast, treatment with either the highly-specific MNK inhibitor MNK-i1 or the eIF4E inhibitor 4EGi-1 did not significantly impact the TOP genes despite both drugs impacting eIF4E. It is possible that mTOR is singular in its effect on their regulation, with minimal effect from convergent pathways. Off-target effects of commonly-used MNK inhibitors in past studies^23^ may overemphasize previous observations to this effect, with off-target effects of 4EGi-1^41^ similarly complicating interpretation of eIF4E targets. 4EGi-1 binds to eIF4E with relatively low affinity, and has been shown to induce ER stress and apoptosis independent of inhibition of cap-dependent translation^30,41^. The substantial concentration of 4EGi-1 required to inhibit eIF4E:G association, combined with off-target and cytotoxic effects manifested through six hours of treatment, potentially incurred TOP gene-preserving compensatory regulation.

In a similar manner, pairwise combinations triggered what we interpreted as compensatory attenuation of their strongest effects. We found a lesser degree of attenuation in the combination of PP242 and BKM120, despite the redundancy of these drugs targeting the same pathway, with a greater portion of their individual targets enhanced by combination treatment, suggesting that inhibition of PI3K with BKM120 does not saturate inhibition of mTOR downstream. These findings highlight the utility of explicitly testing combined perturbations and the need for scalable measurement strategies like riboPLATE-seq.

This study serves as a proof-of-concept for larger-scale perturbation screens of potential translational regulators, demonstrating the sensitivity of riboPLATE-seq to translational regulation at the level of initiation. Here, riboPLATE-seq revealed specific translational targets for kinases consistent with the known structure of their signaling pathways, including the established mechanism by which mTOR controls translation of the TOP motif-containing genes. The method serves as a screening protocol, low in cost, complexity, and specificity, but nonetheless sensitive to changes in gene-specific differences in ribosome association between experimental conditions. Ideally, riboPLATE-seq provides a set of candidates for a given perturbation, which can be examined in more detailed analyses and orthogonally validated in subsequent experiments. The technology described could enable a more comprehensive screen of RNA-binding proteins or kinases, the majority of which remain unstudied at the level of translation. Translational networks inferred from such a dataset could be validated with high-resolution techniques like ribosome profiling and CLIP-seq, and used in further study. We anticipate that the ability to dissect these networks at scale will advance our understanding of translational regulation and the design of specific therapies for diseases involving aberrant translation.

## METHODS

### Tissue Culture and Compound Administration (TS543)

We seeded TS-543 neursopheres (passage #11) on a 96-well plate (Corning, #3799) at a density of 7,500 cells per well (50,000 cells/mL) in 150uL NS-A complete medium (containing 10% v/v NeuroCult NS-A Proliferation Supplement, 20ng/mL EGF, 10ng/mL bFGF, and 2ug/mL heparin) (STEMCELL Technologies #05751). We incubated the plate of cells for 36 hours prior to the start of the experiment at 37°C and 5% CO2 in a tissue culture incubator. We separately prepared stock solutions of PP242 (Tocris, #4257), MNK-i1 (Sigma, **#** 534352), NVP-BKM120 (Selleck, S2247), AZD8055 (Selleck, S1555), and 4EGi-1 (Tocris, #4800) in DMSO vehicle (Sigma, #472301). After dilution with NS-A basal culture medium (without supplement, cytokines, or heparin), we administered the drugs or pure DMSO to the experimental and control wells, respectively, in 1uL doses. Final concentrations were 50nM AZD8055, 625nM PP242, 1μM BKM120, 100nM MNK-i1, and 50μM 4EGi-1, including in pairwise combination-treated samples. Drug treatment proceeded for 6 hours in the tissue culture incubator prior to lysis.

### Cell Lysis (TS543)

Following treatment, we centrifuged the plate of TS-543 for 7 minutes at 1800RPM on a Sorvall Legend XTR at room temperature and removed supernatants by aspiration. Placing the plate on ice, we resuspended the pelleted cells in each well in 30uL of polysome lysis buffer (20mM Tris-HCl, pH=7.4, 250mM NaCl, 15 mM MgCl_2_),0.1mg/mL cycloheximide, 0.5% Triton X-100, 1mM DTT, 0.5U/mL SUPERase-In (ThermoFisher, AM2696), 0.024U/mL TURBO DNase (Life Technologies, AM2222), 1x Protease Inhibitor (Sigma, P8340)), mixed 5 times by pipetting, and rested the plate on ice for 5 minutes. We then centrifuged the plate for 5 minutes at 1400RPM at 4°C to remove bubbles before performing a quick freeze-thaw, placing the plate first in a −80°C freezer and then resting at room temperature for 5 minutes each. Following an additional 10 minutes rest on ice, we viewed the plate under a microscope to check the extent of cell lysis. We then prepared a new 96-well plate containing 3.5uL 2x TCL buffer (Qiagen, #1070498) per well, to which we transferred 3.5uL of lysate (approximately 10% total volume).

### Automated Pan-Ribosome Immunoprecipitation

To the remaining lysate, we added 1 uL of SUPERase-in (ThermoFisher, AM2696) and 1 uL of biotinylated y10b antibody (ThermoFisher, MA516060) to each well, then sealed the plate and allowed it to incubate while gently shaking for 4 hours at 4°C. During this incubation, we washed 500uL of Dynabeads MyOne Streptavidin C1 streptavidin-coated magnetic beads (ThermoFisher, #65001) 3 times with polysome wash buffer (20mM Tris-HCl (pH 7.4), 250mM NaCl, 15mM MgCl2, 1mM DTT, 0.1mg/mL cycloheximide, 0.05% v/v Triton X-100), using 1mL per wash and resuspending in 500uL. We added 5uL of washed beads to each well, then incubated while gently shaking at 4°C for an additional hour. After this short incubation, we placed the plate on a magnet, removed and reserved supernatants, and washed the wells 3 times with 200uL per well of polysome wash buffer supplemented with 1uL/mL SUPERase-in on the Biomek 4000 automated liquid handling system.

Following the final wash, we resuspended the beads in 15uL of ribosome release buffer (20mM Tris-HCl (pH 7.4), 250mM NaCl, 0.5% Triton X-100, 50mM EDTA) per well. During a 15-minute incubation at 4C on a Peltier module, with continuous pipet mixing on the Biomek 4000 in order to maximize elution, we distributed 15uL of 2x TCL buffer to each well of a new 96-well plate. Finally, we replaced the eluted sample plate on the magnet and transferred eluants to the TCL-containing plate.

### Tissue Culture and Cell Lysis for Non-Automated riboPLATE-seq in Wi38

Separately, we seeded WI-38 human fibroblast cells on a 96-well plate at a density of 3,000 cells per well in 60uL cell culture media per well (DMEM (ThermoFisher #11965092) + 10% FBS (ThermoFisher #A3160501)), 36 hours prior to cell lysis. After removing media by aspiration and gently washing wells once with cold PBS supplemented with 0.1mg/mL cycloheximide, we added 30uL cold polysome lysis buffer to each well and mixed by pipetting up and down five times. Additionally, we added 1uL of 1:5000 ERCC spike-in mix 1 (ThermoFisher #4456740) to every other column of wells on the plate. We rested the plate at room temperature for 5 minutes, then ice for 10 minutes more, following which we centrifuged the plate at 1400RPM for 5 minutes to remove bubbles. We then reserved 10uL from each well (33% initial lysate volume) in a second plate for PLATE-seq, and added 10uL 2x TCL buffer (Qiagen) to each well before freezing at −80C.

### Manual Immunoprecipitation of Ribosome Bound RNA (Wi38)

We first added 0.6 uL each of biotinylated antibody y10b (ThermoFisher #) and SUPERase-IN (Life Technologies #) to each well of the remaining lysate, mixed well by pipetting, then sealed the plate and incubated for 4 hours at 4C with gentle shaking. We then added 4uL of streptavidin-coated magnetic beads (DynaBeads MyOne Streptavidin C1, ThermoFisher #65001) to each well, which we had washed 3x and resuspended in polysome wash buffer, mixed carefully, and allowed the plate to incubate for one additional hour at 4C with gentle shaking. After placing samples on a 96-well plate magnet, we washed all wells 3x with polysome wash buffer, and eluted after the final wash by 15 minutes of incubation in 15uL ribosome release buffer. Finally, we removed the supernatant using a 96-well plate magnet, and added 15uL of 2x TCL buffer to each well before freezing the plate.

### Ribosome Profiling and RNA Sequencing

We seeded TS-543 neurospheres in a 6-well plate at a starting density of 50,000 cells/mL in 2 milliliters of NS-A complete medium per well, and allowed the plate to rest for 36 hours. After preparing PP242 solution in DMSO as above, we treated two wells each with 625nM PP242 or DMSO vehicle for 6 hours in the tissue culture incubator. In a separate experiment, we treated two wells each with the same concentrations of PP242 and DMSO for 30 minutes. Following treatment, we transferred samples to 15mL conical vials for centrifugation at 640 RCF for 7 minutes, then removed supernatants and added 400 uL polysome lysis buffer (recipe above). After mixing by rapid pipetting, we transferred samples to 1.8mL microcentrifuge tubes, rested them on ice for 5 minutes, and triturated by 5 passages through a 23-gauge needle. Following a clarifying spin of 11K RCF for 10 minutes at 4C on a benchtop centrifuge, we transferred supernatants to a new set of microcentrifuge tubes and discarded pellets. We prepared ligation-free ribosome profiling and total RNA-seq libraries from the clarified polysome lysates treated for 6 hours following the instructions provided with their respective kits (smarter-seq smRNA-seq kit, Takara-Clontech; NEBnext Ultra-Directional II) augmented with our previously-published ligation-free ribosome profiling protocol^42^. We additionally prepared conventional ribosome profiling and total RNA-sequencing libraries from the samples treated for 30 minutes, using previously-described^38^ modifications to the protocol by Ingolia et al^43^. We sequenced 6 ribosome profiling libraries or up to 12 RNA-seq libraries in one NextSeq 550 high-output 75-cycle kit. Library construction methods and experimental conditions for each sample are presented in Supplementary Table S1. Both PLATE-seq and ligation-free ribosome profiling library preparation protocols are available on our laboratory website (http://www.columbia.edu/~pas2182/index.php/technology.html).

### PLATE-seq Library Preparation and Sequencing

We submitted plates of ribosome-associated and previously reserved total lysate in TCL buffer to the Columbia Genome Center for processing by the previously-described PLATE-seq method of RNA-seq library preparation^6^, which involves poly-A selection of transcripts, incorporation of sequence barcodes in poly(T)-primed reverse transcription, and pooling for subsequent library preparation steps, generating a single 3’-end RNA-seq library from each 96-well plate. We pooled total and ribosome-associated PLATE-seq libraries, sequencing the pooled pairon the Illumina NextSeq 550 with a 75-cycle high-output kit. With paired-end sequencing, the first read corresponds to the 3’ end of a transcript, and the second read contains the barcode identifying the library in which the read was obtained.

### Read Alignment and Data Analysis

With a custom processing pipeline, we first trim reads of trailing polyA sequence and adapters with cutadapt (v 3.5)^44^, then align the whole set of multiplexed reads to the hg38 assembly of the human genome, plus additional sequences corresponding to ERCC spike-in transcripts added for depletion experiments, with STAR^45^ (v 2.7.9a). We then demultiplex the aligned fragments from Read 2 to their original riboPLATE- or PLATE-seq library indices according to their barcodes present in Read 1, as described in the original PLATE-seq paper^6^. We use a similar pipeline to process and align ribosome profiling and RNA sequencing libraries, first trimming polyA tails and adapters with cutadapt, then removing reads that align to the 45S pre-ribosomal RNA and 5S ribosomal RNA with bowtie2^46^ (v 2.2.5) before aligning with STAR. We then use featureCounts^47^ (v 2.0.1) to count the number of fragments aligned to each gene in each library, counting all exon-aligned reads as valid. Barcode sequences for each PLATE-or riboPLATE-seq library generated are available as separate worksheets in Supplementary Table S2.

### Definition of Gene Sets of Interest

As PLATE- and riboPLATE-seq depend on isolation of RNA by poly(T) pulldown, they can only be used to measure polyadenylated transcripts. We first combined two sets of poly(A)- predominant transcripts from HeLa and H9 cells determined in a screen of polyadenylation status across the transcriptome^48^, and removed these genes from consideration in our study to leave only consistently polyadenylated transcripts. We also obtained a set of known 5’ terminal oligopyrimidine motif-containing genes (TOP genes), as well as novel TOP candidates with and without known TOP-containing analogues in mice, from a comprehensive search of transcription start sites^33^.

### Variance-Stabilizing Transformation and Outlier Removal

After subsetting the count matrices for all libraries to remove alignments to non-polyadenylated and spike-in transcripts, we constructed an overall count matrix of all 192 libraries for all 96 samples. We read this matrix into DESeq2 with corresponding column data describing the sample ID, library type (ribo or RNA PLATE-seq), and drug treatment for each library. We then used the variance-stabilizing transform in DESeq2 (version 1.3.4) with default parameters to obtain an approximately homoscedastic, log-scale transformation of raw counts for all libraries. We used this transformed count matrix to perform two-dimensional principal component analyses (PCA) in Python, utilizing the scikit-learn package for analysis and matplotlib for visualization. We limited these analyses to genes determined significant by DESeq2 for differential RA in any drug tested (Benjamini-Hochberg adjusted FDR<0.05; 1813 genes total). For each sample, we computed RA for each gene as the ratio of normalized riboPLATE-seq to PLATE-seq counts; for log-scale transformations such as vst, this corresponds to their difference. With the average RA across all vehicle-treated controls as a reference for baseline RA, we computed the log-fold change from baseline for all genes in each drug-treated sample. We first performed PCA on RA and log-fold change in RA for the full set of samples from the plate, then removed 11/96 samples as PCA outliers (2 DMSO, 2 4EGi-1, 2 BKM120+MNK-i1, 1 AZD8055, 1 PP242, 1 BKM120, 1 PP242+MNK-i1, 1 PP242+BKM120)for subsequent analyses of differential ribosome association with DESeq2. We used the remaining samples to generate final matrices of raw counts and vst-transformed counts for the RNA quality control and principal component analyses in Figures 2 and 3.

### Differential count analysis in DESeq2

We performed differential expression and differential ribosome association analyses using the DESeq2 package in R^49^. We first read the entire matrix of counts across all samples, plus its corresponding column data table describing sample ID, drug treatment, and library type for each sample, into DESeq2. For each condition, we subset the matrix of gene counts to samples corresponding only to that condition and DMSO controls, then analyzed that subset using a likelihood ratio test with the following parameters:

~~~
dds_sub <- DESeq(dds_sub, fitType=‘local’, test=‘LRT’,
full=~condition+type+condition:type, reduced=~condition+type)
~~~

Where `condition` and `type` in the design formulas refer to experimental condition/drug treatment and sequencing library type (riboPLATE or PLATE-seq), respectively. Following each subset DESeq2 analysis, we retrieved results for the interaction term `condition:type`, corresponding to changes in the ratio of riboPLATE-to PLATE-seq counts between conditions, i.e. differential RA:

~~~
res <- results(dds, name=‘condition<X>.typeRIBO’)
~~~

We analyzed ribosome profiling and RNA sequencing data in an identical fashion, comparing PP242 vs DMSO-treated samples at both 30 minutes and 6 hours of treatment to generate two signatures of differential TE.

### Comparison of Sequencing Library Types with Gene Set Enrichment Analysis

We constructed ranked lists for gene set enrichment analysis (GSEA) using the per-gene differential translation efficiencies calculated with ribosome profiling and RNA sequencing data at 30 minutes or 6 hours of PP242 treatment. For each drug tested via riboPLATE-seq, we identified its targets as genes exhibiting significant change in ribosome association (RA) by DESeq2 (Benjamini-Hochberg adjusted FDR< 0.05), split by up- or downregulation (lfcRA>0 / lfcRA<0). We then removed genes from each such that all sets were mutually exclusive, i.e., no two drugs share a common upregulated or downregulated gene. We used the preranked function in the GSEA desktop app to compare the ranked lists for differential TE, using each gene’s log fold change in TE as a ranking metric, against these differential RA-derived gene sets with default parameters and scoring method set to ‘classic’.

### Network Visualization

To create a basic network, we interpreted the genes exhibiting significant reductions in RA under treatment with kinase inhibitors (FDR<0.05) as positive targets of the kinases inhibited. We loaded these gene sets into CytoScape^50^ (v2.9.0) as individual networks for each kinase, merged the three networks, and organized the resulting merged network with the yFiles^51^ Organic automatic layout. We then color-coded the sets of canonical and novel TOP motif-containing genes present in the network, based on lists obtained from Yamashita et al^33^.

### Data Visualization and Code

All code was run in Python 3.9.5 and R 4.0.5. R packages were installed via Bioconductor v 3.12, including DESeq2 v1.34.0 for differential RA and TE analysis and normalization, and BiocParallel v 1.28.0 for use of multicore processors.

Python libraries were installed via Anaconda (v 4.10.3). We generated plots and diagrams using matplotlib (v3.4.3) and Jupyter Notebook (IPython 7.28.0, jupyter_core v4.8.1)^52,53^. Our analyses use NumPy^54^ (v1.21.2) for data manipulation, SciPy^55^ (v1.6.3) for statistical tests, and scikit-learn^56^ (v1.0) for PCA. We additionally generated strip plots and heatmaps using Seaborn^57^ (v0.11.2).

## Acknowledgments

We are grateful to MRC Technology for their gift of the MNK-i1 inhibitor used in this study, as well as Charles Karan and Ronald Realubit of the High-Throughput Screening Facility of the JP Sulzberger Columbia Genome Center for coordinating the automated preparation of our PLATE-seq libraries. NJH was funded by NIH/NINDS F31NS089106. PAS was funded by NIH/NCI R33CA202827.

## Authors Contributions

PAS and NJH initially proposed the riboPLATE-seq protocol, and NJH performed exploratory and foundational work in its development, including the pilot study without automation. JBM and JW developed the workflow for automated ribosome immunoprecipitation. JBM and SDS executed the riboPLATE-seq drug screen, with JBM performing all computational and statistical analyses and SDS culturing cells and preparing lysates. SDS and CG prepared ribosome profiling and RNA-sequencing libraries in this study. JBM and PAS wrote the paper. All authors read and approved the final manuscript prior to submission.

## Data Availability

Sequencing data for this study is available on the Gene Expression Omnibus (GEO) under accession ID GSE139238 [https://www.ncbi.nlm.nih.gov/geo/query/acc.cgi?acc=GSE139238]. Custom scripts used in the analyses performed in this paper, a Jupyter notebook generating our main figures, and the accompanying files necessary for their function are available for download from our laboratory Github (https://github.com/simslab/riboPLATE-seq).

